# Sustained IL-15 response signature predicts RhCMV/SIV vaccine efficacy

**DOI:** 10.1101/2021.01.11.426199

**Authors:** Fredrik Barrenäs, Scott G. Hansen, Lynn Law, Connor Driscoll, Richard R. Green, Elise Smith, Jean Chang, Inah Golez, Taryn Urion, Xinxia Peng, Leanne Whitmore, Daniel Newhouse, Colette M. Hughes, David Morrow, Kurt T. Randall, Andrea N. Selseth, Julia C. Ford, Roxanne M. Gilbride, Bryan E. Randall, Emily Ainslie, Kelli Oswald, Rebecca Shoemaker, Randy Fast, William J. Bosche, Michael K. Axthelm, Yoshinori Fukazawa, George N. Pavlakis, Barbara K. Felber, Slim Fourati, Rafick-Pierre Sekaly, Jeffrey D. Lifson, Jan Komorowski, Ewelina Kosmider, Jason Shao, Wenjun Song, Paul T. Edlefsen, Louis J. Picker, Michael Gale

## Abstract

Simian immunodeficiency virus (SIV) challenge of rhesus macaques (RMs) vaccinated with Rhesus Cytomegalovirus (RhCMV) vectors expressing SIV proteins (RhCMV/SIV) results in a binary outcome: stringent control and subsequent clearance of highly pathogenic SIV in ~55% of vaccinated RMs with no protection in the remaining 45%. Although previous work suggests that unconventionally restricted, SIV-specific, effector-memory (EM)-biased CD8^+^ T cell responses are necessary for efficacy, the magnitude of these responses does not predict efficacy, and the basis of protection vs. non-protection in RhCMV/SIV vector-vaccinated RMs has not been elucidated. Here, we report that RhCMV/SIV vector administration strikingly alters the whole blood transcriptome of vaccinated RMs, with the sustained induction of specific immune-related pathways, including non-canonical T cell receptor (TCR), toll-lie receptor (TLR), inflammasome/cell death, and interleukin-15 (IL-15) signaling, significantly predicting protection. The IL-15 gene expression signature was further evaluated in an independent RM IL-15 treatment cohort, revealing that in whole blood the response to IL-15 is inclusive of innate and adaptive immune gene expression networks that link with RhCMV/SIV vaccine efficacy. We also show that this IL-15 response signature similarly tracks with vaccine protection in an independent RhCMV/SIV vaccination/SIV challenge RM validation cohort. Thus, the RhCMV/SIV vaccine imparts a coordinated and persistent induction of innate and adaptive immune pathways featuring IL-15, a known regulator of CD8^+^ T cell function, that enable vaccine-elicited CD8^+^ T cells to mediate protection against highly pathogenic SIV challenge.

**Author Summary:** SIV insert-expressing vaccine vectors based on strain 68-1 RhCMV elicit robust, highly effector-memory-biased T cell responses that are associated with an unprecedented level of SIV control after challenge (replication arrest leading to clearance) in slightly over half of vaccinated monkeys. Since efficacy is not predicted by standard measures of immunogenicity, we used functional genomics analysis of RhCMV/SIV vaccinated monkeys with known challenge outcomes to identify immune correlates of protection. We found that arrest of viral replication after challenge significantly correlates with a vaccine-induced response to IL-15 that includes modulation of T cell, inflammation, TLR signaling, and cell death programming. These data suggest that RhCMV/SIV efficacy is not based on chance, but rather, results from a coordinated and sustained vaccine-mediated induction of innate and adaptive immune pathways featuring IL-15, a known regulator of CD8^+^ effector-memory T cell function, that enable vaccine-elicited CD8^+^ T cells to mediate efficacy.

## Introduction

Human immunodeficiency virus (HIV) infection continues to be a major global health problem, with approximately 38 million people worldwide currently living with HIV. Despite the decline in new infections and the remarkable success of current antiretroviral therapy (ART) at suppressing viral load in people undergoing treatment, there were 1.7 million new HIV infections in 2019 and nearly 700,000 AIDS-related deaths (1). Thus, the need for a vaccine to protect against HIV infection remains high, underscoring the continued essential role of RM models of SIV infection for developing and testing HIV vaccine concepts. In this regard, the highly pathogenic SIVmac251 swarm and SIVmac239 clones have been especially high bars for prophylactic vaccine efficacy with few concepts reproducibly showing sufficient efficacy against these highly pathogenic SIVs for clinical translation (2–4).

Among these is the T cell response-targeted SIV vaccine that uses vaccine vectors based on RhCMV, which elicit and indefinitely maintain high frequency, circulating and tissue-based, effector memory (EM)-differentiated SIV-specific T cell responses (2, 5–7). RhCMV vectors were designed to provide for an immediate effector T cell intercept of immune-evasive pathogens, so as to implement anti-pathogen immune activity prior to development of effective immune evasion (8). In contrast to conventional T cell-targeted prime-boost vaccines against SIV, which elicit responses that at best suppress, but never clear SIV infection, RhCMV/SIV vector-elicited immune responses have the ability to mediate complete arrest of mucosal-acquired SIV infection prior to establishment of a long-lived SIV reservoir, ultimately resulting in progressive decline of SIV-infected cells in tissues to the point where replication-competent SIV can no longer be detected by the most sensitive assays (9–11). This remarkable “control and clear” efficacy is not, however, universal, but rather occurs in ~55% of vaccinated RM across multiple studies, with the SIV infections in the ~45% of vaccinated, non-protected RMs showing indistinguishable viral dynamics from the unvaccinated controls (9, 10, 12).

Importantly, the RhCMV/SIV vaccine vectors manifesting this efficacy are based on the 68-1 strain of RhCMV, which developed a unique genetic rearrangement during *in vitro* passage that abrogated the function of eight distinct immunomodulatory gene products encoded in two RhCMV genomic regions (Rh157.5/.4 and Rh158-161). These gene modifications had the remarkable effect of programming both the virus- and SIV insert-specific CD8^+^ T cells induced by this vector to recognize epitopes restricted by either MHC-E or MHC-II, but not classical MHC-Ia (13–15). Differential repair of these genes reverts epitope restriction of vector-elicited CD8^+^ T cells to MHC-Ia or MHC-Ia mixed with MHC-II, whereas another mutation of 68-1 RhCMV (deletion of Rh67) results in exclusively MHC-II-restricted CD8^+^ T cells (15, 16). Although the MHC-Ia- and/or MHC-II-restricted CD8^+^ T cell responses elicited by these other RhCMV/SIV vectors manifest similar phenotypic and functional characteristics, only the original 68-1 vector shows efficacy against SIV (15, 16), strongly implicated the MHC-E-restricted, SIV-specific CD8^+^ T cells as the active adaptive component of this non-antibody-inducing vaccine. However, to date, no quantitative parameter of the 68-1 RhCMV/SIV-induced T cell responses has consistently correlated with the binary outcome (protection vs. non-protection) after SIV challenge (9, 10, 12). These observations raise the question of whether protection mediated by MHC-E-restricted CD8^+^ T cells is stochastic – i.e., based on the chance, relative distribution of infection trajectory vs. effector cell distribution in an early, critical time window after viral entry – or based on more complex characteristics of the vaccine-elicited immune response.

Here, we performed functional genomics analyses including bioinformatics linear modeling of the RM whole blood transcriptome before and longitudinally after 68-1 RhCMV/SIV vaccination, asking whether a vaccine-induced, transcriptomic response could predict RMs that were or were not subsequently protected after SIV challenge. These analyses revealed a complex transcriptomic response to vaccination that included modulation of specific immunoregulatory gene pathways linked with IL-15 signaling. Indeed, as assessed the *in vivo* whole blood response to IL-15 to reveal that IL-15 signaling directs a remarkable breadth of gene expression networks linked with innate and adaptive immune programming that underlie RhCMV/SIV vaccine efficacy. Our study defines the gene and gene network correlates mediating vaccine efficacy and protection. These findings that suggest RhCMV/SIV vaccine protection is not stochastic, but rather is dependent on vaccine-induced specific innate and adaptive immune responses.

## Results

### Characterization of a RhCMV/SIV vector-vaccinated RM cohort with known challenge outcome

Two cohorts of male RM (n=15 each) were administered a vaccine composed of three 68-1 RhCMV/SIV vectors individually expressing SIV Gag, SIV Rev/Tat/Nef and SIV 5’-Pol, one group via a subcutaneous (subQ) route and the other via an oral route. Each RM was vaccinated twice, receiving a week (wk) 0 prime (Pr) and wk18 homologous boost (Bo) (**Fig. 1A**). The development of SIV-specific T cells was monitored by intracellular cytokine staining (ICS), with immunogenicity in both the subQ and oral vaccine groups showing robust induction of SIV Gag-, Rev/Tat/Nef-, and Pol-specific CD4^+^ and CD8^+^ T cells (**Figs. S1A,B)**, including CD8^+^ T cell responses to previously characterized MHC-E- and MHC-II-restricted SIV Gag supertopes – indicating, as expected, the induction of the unconventionally restricted CD8^+^ T cells; **Fig. S1C**) (13, 14). These T cell responses, which were maintained through the time of challenge, manifested a striking EM bias in phenotype and function, similar to our previous analysis of T cell responses elicited by these vectors (5, 9, 10, 12) (**Fig. S1D-E**). Of note, there were no significant differences in these immunogenicity parameters in RM given the vaccines via subQ vs. oral route. At wk91 after initial vaccination, these vaccinated RM (and 15 unvaccinated controls) were subjected to repeated limiting dose, intrarectal challenge until establishment of infection “take” by detection of *de novo* T cell responses to SIVvif (an SIV Ag not included in the vaccine; **Fig. 1B**), as previously described (5, 9, 10, 12). Analysis of plasma viral load in RM with such infection “take” showed stringent control of viral replication in 8 and 9 of 15 RMs from the subQ and oral groups (protected RMs), respectively, with no “protected” RMs in the unvaccinated control group (**Fig. 1C**). Analysis of tissue cell-associated viral load confirmed that protected RMs were indeed infected by SIV (**Fig. 1D**), demonstrating that the absence of viremia in these RMs reflected immune-mediated arrest of SIV replication. Pre-specified statistical analyses of the T cell responses measured by ICS across the two vaccine cohorts found no association between magnitude and phenotype of the SIV-specific responses and protection from progressive SIV infection across the two vaccinated RM cohorts (**Figs. S2A-E)**.

**Figure 1.**
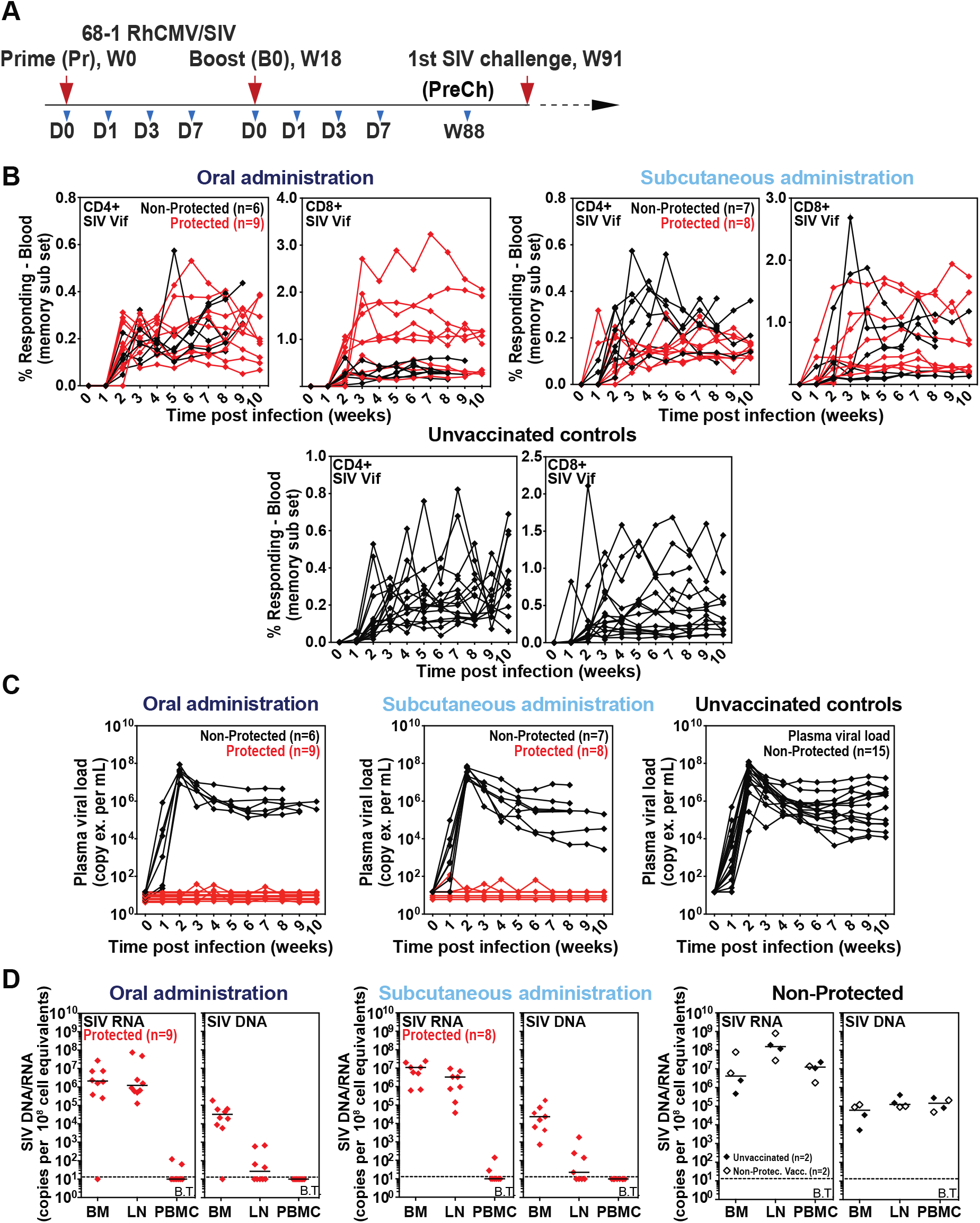
Virologic and immunologic responses in Rh/CMV vaccination and SIV challenge. (**A**) Schematic of the vaccine phase of the two cohorts of RM (n = 15 each) administered the 68-1 RhCMV/SIV vector set by either subcutaneous or oral routes at wk0 Pr and wk18 Bo, indicating time points for which whole blood samples were collected for RNAseq analysis. Repeated limiting dose SIV_mac239_ challenge was initiated at wk91. (**B,C)** Assessment of the outcome of effective challenge by longitudinal analysis of the *de novo* development of SIV Vif-specific CD4^+^ and CD8^+^ T cell responses (**B**) and plasma viral load (**C**). RM were challenged until the onset of any above-threshold SIV Vif-specific T cell response, with the SIV dose administered 2 or 3 weeks prior to this response detection considered the infecting challenge (week 0). RM with sustained viremia were considered not protected; RM with no or transient viremia were considered protected (9, 10, 12). (**D)** Bone marrow (BM), peripheral lymph node (LN) and peripheral blood mononuclear cell (PBMC) samples from all vaccine-protected RM and representative non-protected or unvaccinated control RM, collected from between day 28 and day 56 post-SIV infection, were analyzed by nested, quantitative PCR/RT-PCR for cell-associated SIV DNA and RNA. The horizontal line indicates the threshold of detection (B.T. = below threshold) with data points below this line reflecting no positive reactions across all replicates. Above threshold cell-associated SIV RNA was detected in LN and BM of all protected RM, confirming SIV infection take.

### Analysis of the whole blood transcriptome induced by subQ and oral RhCMV/SIV vaccination

To determine whether other parameters of the vaccine response might serve to differentially program protective immunity across the outcome groups, we performed longitudinal global transcriptomic profiling of mRNA expression in whole blood samples collected prior to Pr and Bo RhCMV/SIV vaccinations (d0 of wk0 and wk18, respectively], at d1, d3 and d7 following Pr and Bo, and at wk88, 3 wks prior to initiating SIV challenge (see **Fig. 1A**). We first conducted analyses comparing baseline gene expression profiles across subQ and oral cohorts and found that pre-vaccination signatures of individual RMs were similar to one another across cohorts (**Figs. 2A, B**). We next evaluated the gene expression responses to vaccination by performing a principal component analysis (PCA) on the per-timepoint mean log2 fold-change (FC) values, using all 12,734 expressed genes and averaged over the RMs within each treatment and outcome group (**Fig. 2C**). PC1 and PC2 constituted the majority of the post-vaccination gene expression variability that segregated the animals into protected and non-protected outcome groups.

**Figure 2.**
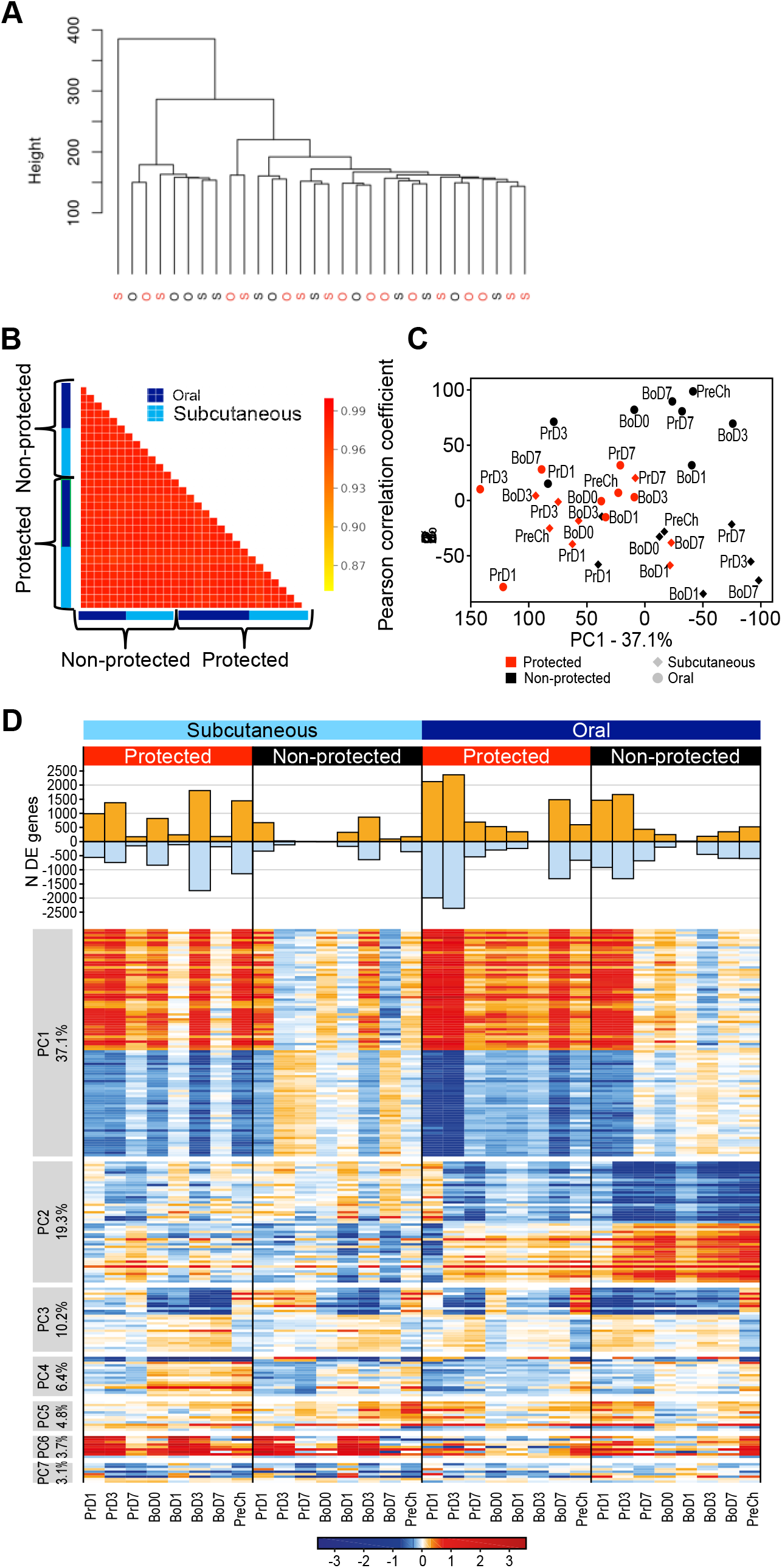
Identification of DE genes after RhCMV/SIV vaccination in protected vs. non-protected RM. Figure S3. Wk0, d0 signature comparison. **(A, B)** Week 0, d0 signature comparisons. (**A**) Ward hierarchical clustering of animals by their log count per million (CPM) values at baseline was performed to evaluate initial similarity between animals. O and S indicate animals from oral or subQ vaccine administration, respectively. Red: protected, black: non-protected. (**B)** Pearson correlation matrix of animals based on their log CPM values at baseline to evaluate initial similarity. Animals are grouped by protection status and oral (green) or subQ (blue) vaccine administration. (**C**) PCA of the per-time point mean log2 fold-change (FC) values, using all expressed genes and averaged over the RMs within each treatment and outcome group, showing the mean log2(FC) of all expressed genes per time point in oral (circles) and subcutaneous (diamonds) for protected (red) and non-protected (black) RMs. (**D)** Upper: number of DE genes per time point in each group. Lower: Heatmap showing genes most associated with each PC, for the first 7 principal components. Percent variance explained by each PC is shown at left. Prime, boost, and pre-challenge time points are shown at bottom.

To further evaluate these differences, we defined the set of genes with statistically significant differential expression (DE) at any post-vaccination time point compared to baseline in any of the four RM subgroups defined by administration route and protection outcome, using bioinformatics analyses and linear modeling (**Fig. 2D; Table S1**). These analyses suggested that DE genes associated with variation in PC1 are relevant for establishing a vaccine protective signature. PC1 DE gene expression changes occurred rapidly after vaccination (d1 in both cohorts), with highest numbers and absolute levels of gene expression change from baseline found in the oral cohort 3d after Pr. The expression of these DE genes was variably reduced thereafter, but were relatively maintained in the protected RMs, with the expression level change increasing to a greater degree in the protected group following vaccine boost. Importantly, absolute gene expression changes of the PC1 DE genes persisted through the pre-challenge (PreCh) time point in the protected RMs, whereas non-protected RMs failed to maintain or re-establish this signature by the PreCh time point.

### Gene network correlates of protection

To identify gene expression correlates of protection and their functional regulatory networks, we determined the DE genes showing significant differential expression between protected and non-protected groups across the time course, designated as DDE genes. The 2,272 DDE genes identified (**Table S2**) capture the variable signature between protected and non-protected outcome groups (**Fig. 3A)**. Using permutation testing we verified that the magnitude of differential expression between protection outcomes was significant (P = 0.011), thus linking DDE gene expression with vaccine protection. To identify specific response pathways and networks among the DDE genes that define vaccine protection, we first performed clustering analysis and found four major clusters containing either mostly induced/up-regulated genes (Clusters 1 and 2) or suppressed/down-regulated genes (Clusters 3 and 4) in the protected RM across both vaccine groups (**Fig. 3A**). The vaccine-induced gene clusters were associated mainly with innate immune pathways spanning TLR signaling and cytokine response pathways, inflammasome/cell death signaling, and included a set of T cell/TCR signaling genes (**Fig. 3B**). Pathway analyses of DDE gene network linkage identified death receptor signaling/inflammasome network interactions with a TLR network and genes involved in TCR signaling marked through previously defined interactions of *PPP3CA*/calcineurin and *PTPN11*/SHP1 phosphatases (17–20), and phosphatidylinositol responsive kinase (PI3K) subunits (*PI3KR1* and *PIK3CB)(21)*, each network significantly associated with vaccine protection (**Fig. 3C**). Notably, genes encoding essential factors of T helper-1 polarization and immune activation, including the T-bet transcription factor (*TBX21*), IL-2 and IL-12 receptor subunits, and the Zeta associated protein kinase 70 (*ZAP70*)(22–24) were suppressed or “down-regulated” concomitant with up-regulation of *PTPN11, PPP3CA* and indolamine dioxegenase 1 (*IDO1*), genes known to impose an immune regulatory phenotype(25, 26), thus defining a non-canonical T cell activation/TCR signaling signature in the blood of protected RMs.

**Figure 3.**
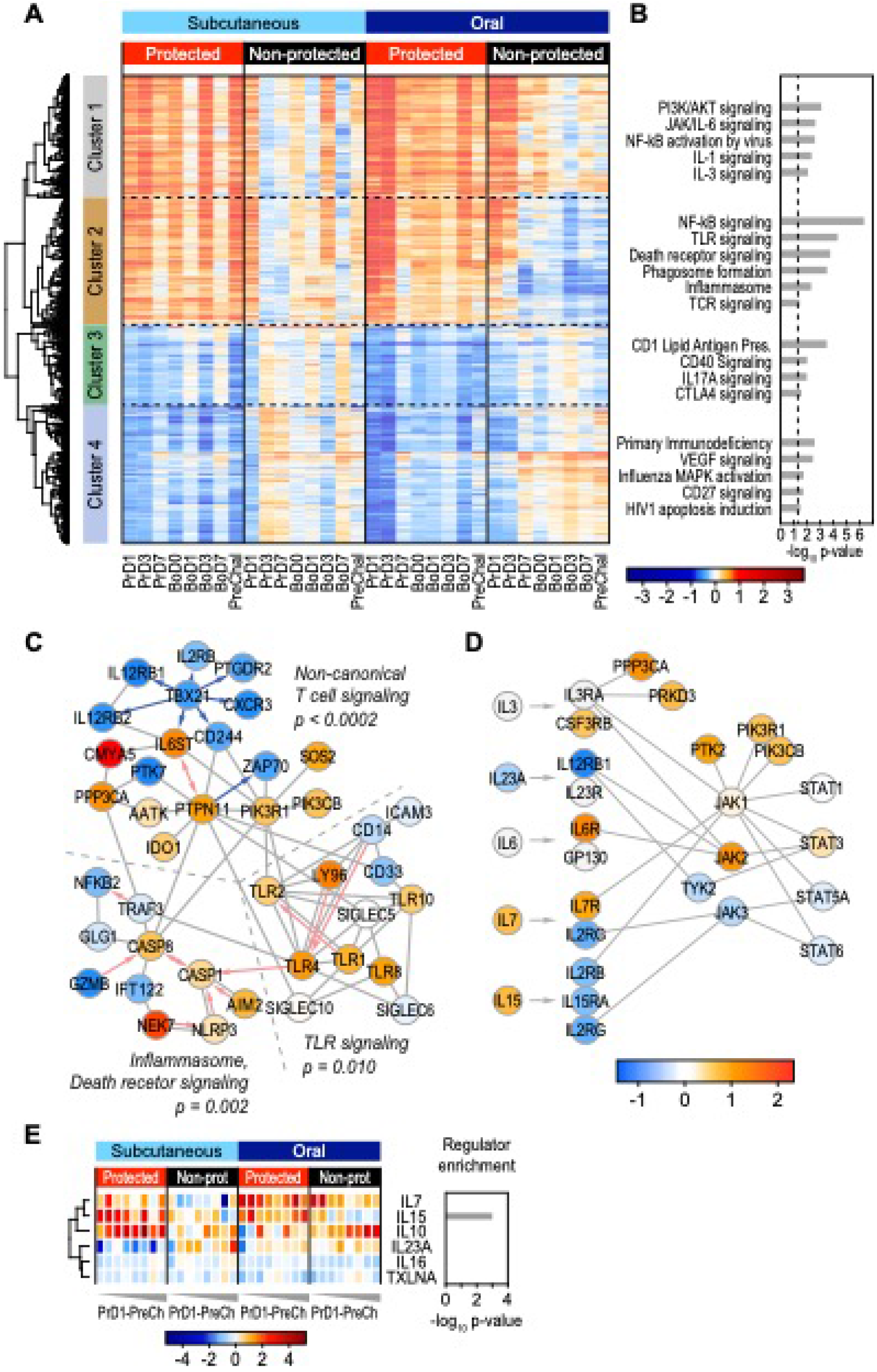
Gene and pathway correlates of protection. (**A**) Heatmap showing all DDE genes. Four clusters were defined using hierarchical clustering. (**B**) Ingenuity pathway analysis of the four major DDE gene clusters. (**C**) Network of direct physical interactions between major enriched immune pathways with red and blue arrows indicating activating and inhibitory interactions, respectively. P-values for the association of each pathway with vaccine protection are based on permutation testing. (**D**) Network overview of JAK-STAT signaling in enriched interleukin pathways. (**E**) Heatmap of gene expression changes for expressed interleukin genes (**left**), and their enrichment as upstream regulators of DDE genes using Ingenuity analysis (**right**).

We also found that JAK-STAT cytokine signaling was enriched within the DDE gene signature in the absence of typical type I interferon stimulated genes (ISGs), suggesting different interleukin signaling programs (**Fig. 3D**). We therefore interrogated our gene expression data sets for expression levels of all interleukins, identifying only IL-7, IL-10, TXLNA/IL-14, IL-15, and IL-23A as being expressed at one or more time points within the vaccine cohorts. Among these, IL-15 expression was enriched in protected RMs and was identified as a significant upstream regulator of the vaccine protection-associated DDE signature (**Fig. 3E**). We found that the increased upregulation of IL-15 expression in protected, compared to nonprotected, RMs was accompanied by downregulation of IL-15 receptor subunits and their JAK-STAT signaling components relative to the other interleukins. Downregulation of cytokine receptor expression is a specific marker of active cytokine signaling (27), suggesting that an active IL-15 cytokine signaling pathway is a component of the DDE protection signature (see **Fig. 3D**).

### Protection signature links with IL-15 signaling

IL-15 plays a major role in cellular immune programming, supporting memory T cell and NK cell activation, homing, homeostasis, both effector differentiation and function, and in particular controlling the activity of circulating and tissue resident CD8^+^ EM T cells (28–32). IL-15 activates signaling through the β chain and common γ chain heterodimer of the IL-2 receptor, either as a soluble heterodimer with the α chain of the IL-15 receptor (IL-15Rα) or through trans-presentation by cells expressing IL-15Rα(33–35). Mimicking this process, recombinant, purified heterodimeric IL-15/IL-15Rα (rRh-Het-IL-15) is a potent immune therapeutic to induce IL-15 signaling (36–38). To further define the whole blood gene expression signature directed by IL-15 *in vivo* and to evaluate the breadth of the IL-15 response in 68-1 RhCMV/SIV vaccinated RMs, we conducted transcriptomic analysis on whole blood samples from a separate cohort of five unvaccinated RM treated with rRh-Het-IL-15 in a dose-escalating fashion (5, 10, and 15 μg/kg at d0, d3 and d7 respectively) with blood collection through d29 (**Fig. 4A**). We identified DE genes responding to rRh-Het-IL-15 treatment *in vivo* across this 29d time course (**Table S3**) in which IL-15 rapidly altered gene expression within one day of administration followed by a homeostatic reset of expression levels two days later (**Fig. 4B, left panels**). We used the d1 post IL-15 administration DE genes to interrogate the DDE signature of the vaccinated RMs for evidence of an embedded IL-15 response. This d1 DE gene set was selected to reflect the direct response of the RMs to IL-15 prior to the onset of homeostatic regulation. We identified multiple co-expression clusters within the intersecting gene set of 256 genes (**Fig. 4B, Table S4**). Pathway analyses of each cluster revealed an overlap of the response to rRh-Het-IL-15 with several of the immunological pathways identified in the DDE analysis, including up-regulated pathways (cluster A) of PI3K signaling, NF-kB-driven inflammatory genes, death receptor signaling, and T cell signaling modules (**Fig. 4C**). These networks link to interferon regulatory factor (IRF) and STAT transcription factors as major upstream regulators responding to IL-15 signaling. Moreover, we identified acutely down-regulated IL-15 response genes that were also components of the DDE protection signature (see **Fig. 4C**, cluster B). Notably, among genes showing acute downregulation by rRh-Het-IL-15 treatment were pathways regulated by *TBX21*/Tbet, STAT6 and other upstream regulators, consistent with the non-canonical T cell and cytokine signaling DDE signatures (see **Figs. 3C,D**).

**Figure 4.**
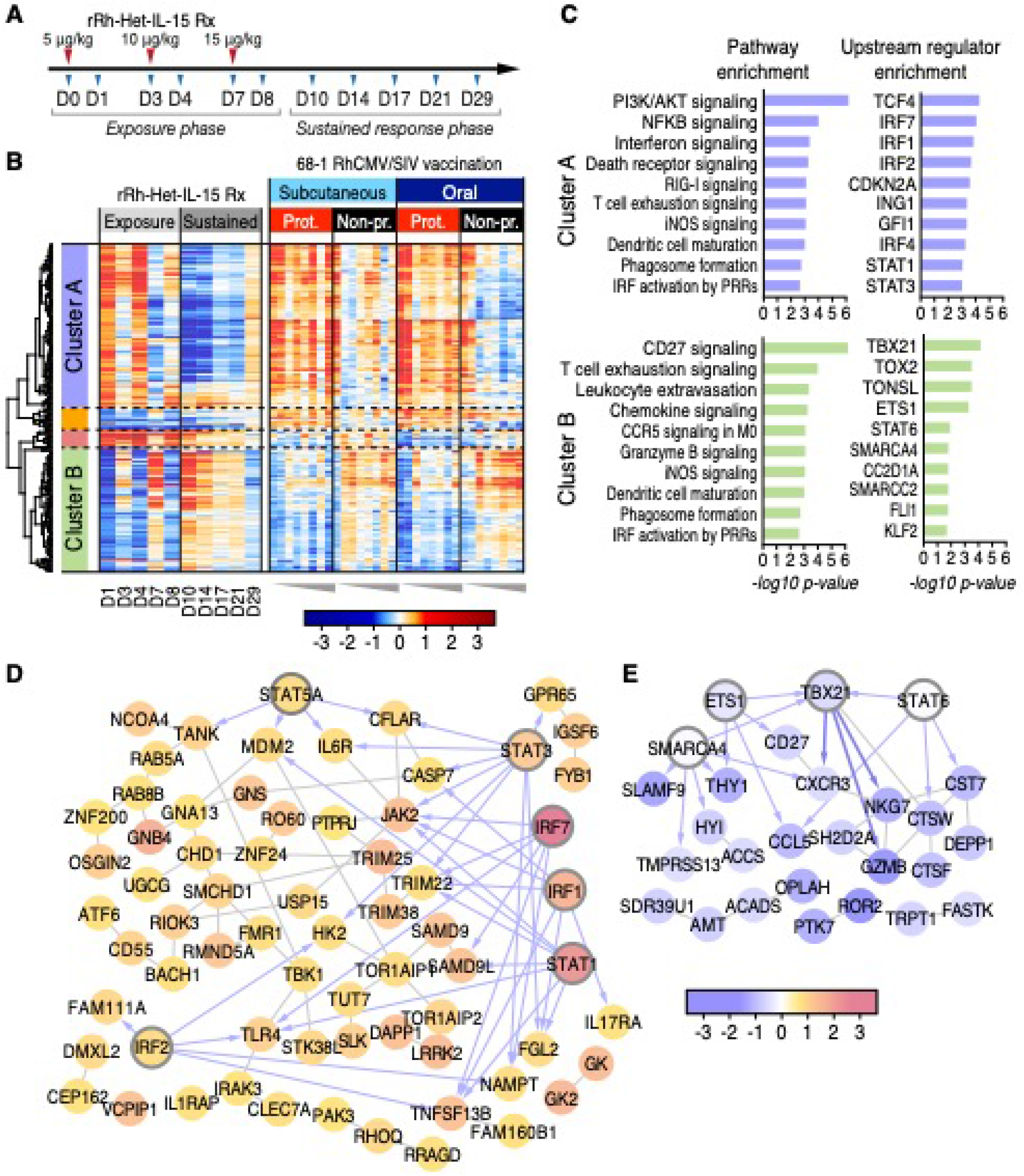
The IL-15 response links with correlates of vaccine protection. (**A**) Study design – RM treatment with rRh-Het-IL-15. (**B**) Heatmap of DDE gene correlates of protection regulated by IL-15 (IL-15 DE genes on d1 post-administration). (**C**) Ingenuity pathway and upstream regulator enrichment analyses for gene cluster A and B from panel B. **(D, E)***De novo* network of genes from Cluster 1 (D), and (E) Cluster 2 were constructed in GeneMania. Transcription factor nodes are indicated by thick borders with connections shown as gray lines and blue arrows, showing co-expression interactions (GeneMania) and direct transcription factor-target interactions (Ingenuity), respectively.

To identify interaction between IL-15 regulated genes within the DDE protective signature, we built *de novo* gene interaction networks using GeneMania (39), basing network construction on genes within each cluster of our intersecting dataset. Among genes up-regulated by rRh-Het-IL-15 treatment within the DDE vaccine protection signature, we identified multiple pathways involved in immune activation including TLR signaling, innate immune activation, and death receptor signaling, each linked to specific transcription factor nodes (IRF1, IRF2, IRF7, STAT1, 3, and 5). As these IRFs and STATs are also prominent ISGs (40), these results together reveal a remarkable breadth of signaling crosstalk in immune programming wherein IL-15 signaling intersects both innate and adaptive immune pathways in building the DDE protection signature, likely reflecting a cascade of direct and indirect IL-15 signaling actions (**Figs. 4D,E**). Of note, we identified a linkage of the acutely IL-15 down-regulated genes with specific transcription factors of the non-canonical T cell activation TCR signaling DDE module described above (**Fig. 4E**). Most importantly, the acute IL-15 response gene set (all IL-15 DE genes at d1) was significantly linked with vaccine protection, using permutation testing methods similar to how we evaluated the DDE signature (P = 0.002). These analyses identify IL-15 as a major regulator of the DDE signature underlying RhCMV/SIV vaccination outcome and demonstrate linkage of IL-15 responsive genes with vaccine efficacy.

### Conserved IL-15 response links with vaccine protection in validation cohort analysis

We next evaluated the whole blood gene expression signature in an independent validation cohort of 15 RM that had been subQ vaccinated with a combination vaccine including the same 68-1 RhCMV/SIV vaccine set used in the RM described above and a variant 68-1.2 RhCMV/SIV vaccine set with the same SIV inserts (**Table S5)**. The 68-1.2 RhCMV/SIV vector is repaired for pentameric complex expression and therefore is programmed to elicit MHC-Ia-restricted CD8^+^ T cell responses, resulting in these vaccinated RM having both conventionally (MHC-Ia) and unconventionally (MHC-E and MHC-II) restricted SIV-specific CD8^+^ T cell responses (15). 68-1.2 RhCMV/SIV vectors are not protective, but they do not abrogate 68-1 RhCMV/SIV vector-mediated protection, and 6 of the 15 vaccinated RM in this validation cohort manifested stringent viral control after SIV challenge (15). Although the 68-1+68-1.2 combination vaccine is quite different than the 68-1-only vaccine, the protection phenotype in RM (e.g., viral replication-arrest) was identical to our 68-1-only vaccinated cohorts, suggesting that protection-critical signaling such as the IL-15 pathway described in **Fig. 4**, might be preserved in protected animals receiving this combination vaccine. The validation 68-1+68-1.2 cohort was studied in parallel with the subQ and oral, 68-1 vaccinated cohorts, and sample collection, processing and RNAseq analysis were performed identically. Comparison of the post-vaccination change-from-baseline expression pattern of the IL-15-regulated genes shown in **Fig. 4B**(intersection of DDE and day 1 rRh-Het-IL-15 DE) in protected vs. non-protected RM in the validation cohort and the SubQ 68-1 vaccinated cohort (also n = 15) revealed a very similar pattern with protected RM in both cohorts showing a more pronounced and durable IL-15 response to vaccination than the non-protected RMs (**Fig. 5A; Tables S4, S6**). In keeping with this outcome, the vaccine response in protected animals in both cohorts showed a higher correlation with the IL-15 response than in non-protected animals (**Fig. 5B**).

**Figure 5.**
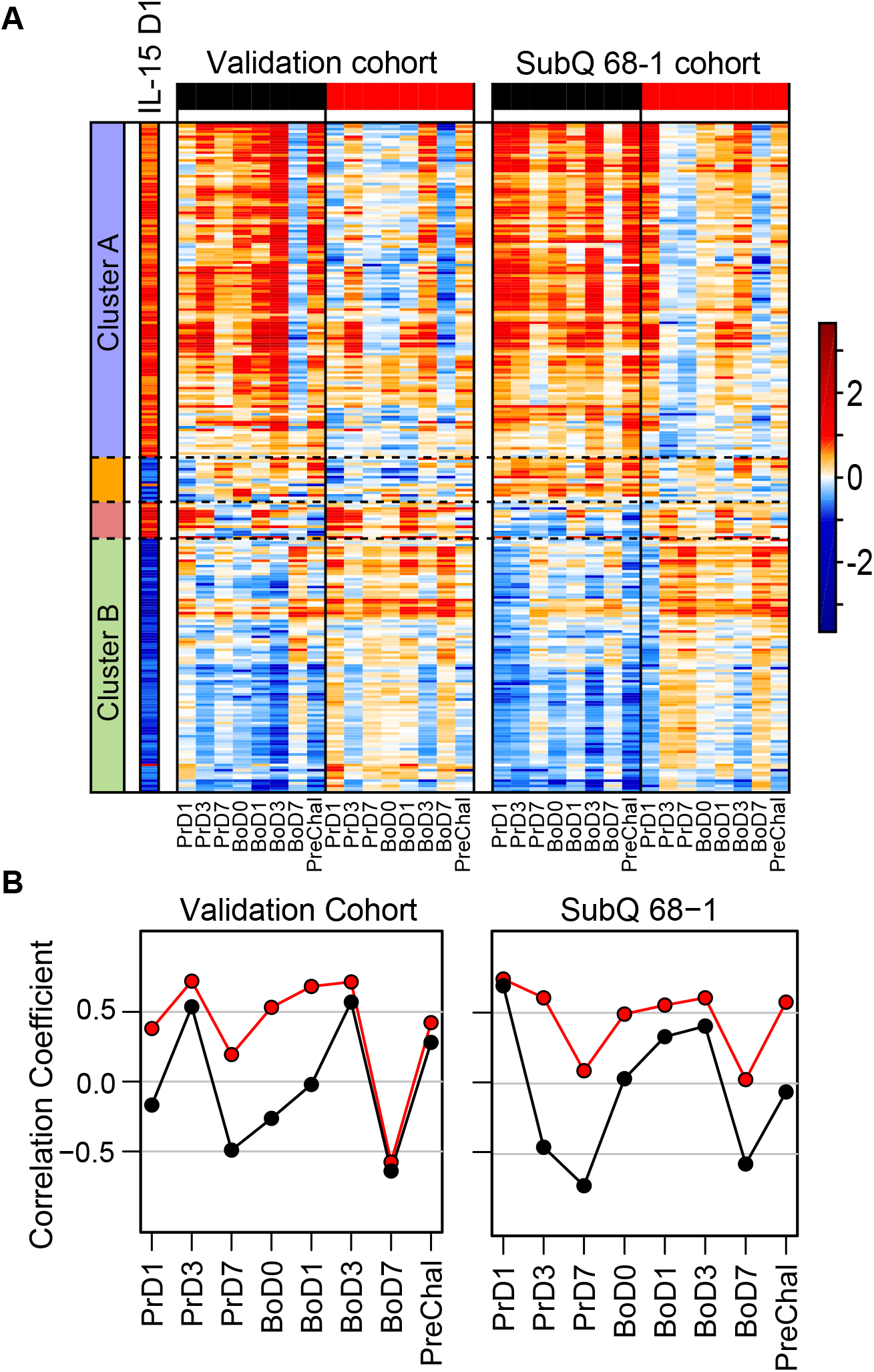
Validation of the IL-15 response signature of protection. (**A**) Heatmaps comparing the IL-15 response signature (overlap of DDE and day 1 IL-15 DE genes as shown in **Fig. 4 B**) in the validation cohort (left panel; 68-1 + 68-1.2 RhCMV/SIV vaccinated; 6 of 15 protected) vs. the original subQ 68-1 RhCMV/SIV vaccinated cohort (right panel; 8 of 15 protected). P-values for the association of the IL-15 response signature with vaccine protection are 0.074 and 0.030 for the validation and SubQ cohorts, respectively (based on permutation testing). (**B**) Pearson correlation coefficients between the IL-15 response signature log2(FC) after one day of IL-15 administration and the group average log2(FC) at each time point from the DE analysis of the validation cohort (**left**) and the 68-1 subQ vaccinated cohort (**right**). Red and black points indicate correlation with values in protected and non-protected RMs, respectively.

## Discussion

Our study reveals an IL-15 response gene signature underlying Rh/CMV-SIV vaccine protection in RM. This signature encompasses functional pathways/modules consisting of non-canonical T cell signaling (defined in this study), TLR signaling, and inflammasome/cell death signaling that correlate with vaccine protection. The magnitude and persistence of gene expression changes following vaccination are significantly greater and sustained throughout the study time course in RM destined for protection after challenge with the IL-15 response signature being a predictor of a vaccine response that links with protection across vaccination cohorts.

IL-15 is produced by myeloid cells, including dendritic cells and monocyte/macrophages, and acts on T cells and NK cells to activate/enhance effector response and cell homing actions (28–32). Since adaptive (SIV-specific) cellular immunity is required for RhCMV/SIV-based vaccine efficacy (9), the most likely targets of this cytokine in our protected RM are the unconventionally MHC-E-restricted SIV-specific CD8^+^ T cells that are both uniquely elicited by this vaccine and associated with efficacy (15, 16). These observations support a model in which vaccine-induced IL-15 production from myeloid-derived cells acts on MHC-E-restricted, SIV-specific CD8^+^ T cells to facilitate the effector activities that result in systemic arrest of viral replication in vaccinated, SIV-challenged RM. Thus, we postulate “tuning” by persistent induction of IL-15 and associated innate immune and immunoregulatory pathways may be required for the MHC-E-restricted, SIV-specific CD8^+^ T cells to mediate arrest of viral replication.

Our study also examined the whole blood transcriptional response to administration of bioactive IL-15. Our functional genomics analyses of the whole blood response shows that IL-15 induces a remarkable breadth of innate and adaptive immune gene expression including engagement of genes and gene networks linked to lymphocyte activation, migration and homing, and innate immunity including pathogen recognition receptor signaling, interferon regulatory factor signaling, and type I interferon actions (see **Fig 4**). Our study design of serial IL-15 and dose-escalating administration shows that IL-15 directs a rapid alteration of the blood transcriptome within 1 day to both induce and suppress specific gene expression from pretreatment baseline levels with similarly rapid homoeostatic “resetting” of the blood gene expression profile two-days later. Moreover, repeated IL-15 dosing and dose escalation generated a decreased response by day 8 post-initial treatment followed by homeostatic regulation of gene expression through the remaining 21 days of the time course. These results show that the blood response wanes and resets after IL-15 withdrawal. By comparing the whole blood response to IL-15 with the RhCMV/SIV vaccine protection signature we were able to identify the IL-15 response genes of vaccine protection and show that the persistence of this expression signature tracks with vaccine efficacy. The persistence of the IL-15 signature in the vaccinated and protected animals suggests that RhCMV-based vaccination induced IL-15 production by one or more cell and/or tissue types *in vivo*, and that protection from SIV infection occurs when this response persists at least to the time of a virus challenge. As RMs receiving IL-15 became refractory or exhibited reduced response to high dose IL-15 after serial administration, the sustained IL-15 signature in the vaccinated protected animals supports the notion that low, persistent IL-15 production links with RhCMV-SIV vaccine efficacy. Thus, strategies to deliver or produce a low, sustained production and response to IL-15 may serve as effective adjuvant approaches to enhance vaccine efficacy.

The basis for the differential induction and maintenance of the protection-associated signaling signature among individual RM is not clear. It is, however, unlikely that differences among RM in vector spread and persistence account for this heterogeneity, as typical protection is observed with pp71-deleted RhCMV/SIV vectors that manifest ~1000-fold reduction in *in vivo* spread compared to the vectors used in this study (10). More likely, host differences in regulation and counter-regulation of these pathways including the production and response of IL-15 underlie protection vs. non-protection. This protective signature is an important correlate of RhCMV/SIV vaccine efficacy in RMs, and thus will be crucial for guiding clinical development of an HCMV/HIV vaccine to recapitulate RhCMV/SIV vector efficacy in humans.

## Materials and methods

### Rhesus macaques

The experiments reported in this study used a total of 65 purpose-bred male and female RM (*M. mulatta*) of Indian genetic background, including 15 RM assigned to each of four vaccine groups (oral 68-1 vaccination, subQ 68-1 vaccination, an unvaccinated control group and a subQ 68-1 + 68-1.2 vaccinated test group), and 5 RM administered rRh-Het-IL-15 for the IL-15 blood signature assessment. The unvaccinated and subQ vaccinated RM cohorts are also reported in Malouli, *et al* (15). At assignment, all study RM were free of cercopithecine herpesvirus 1, D-type simian retrovirus, simian T-lymphotrophic virus type 1, and *Mycobacterium tuberculosis*, but were naturally RhCMV-infected. All study RM were housed at the Oregon National Primate Research Center (ONPRC) in Animal Biosafety level 2 (vaccine phase) and level 2+ (challenge phase) rooms with autonomously controlled temperature, humidity, and lighting. Study RM were both single- and pair-cage housed. Animals were only paired with one another during the vaccine phase if they belonged to the same vaccination group. All RM were single cage-housed during the challenge phase due to the infectious nature of the study. Regardless of their pairing, all animals had visual, auditory and olfactory contact with other animals. Single cage-housed RM received an enhanced enrichment plan that was designed and overseen by RM behavior specialists. RM were fed commercially prepared primate chow twice daily and received supplemental fresh fruit or vegetables daily. Fresh, potable water was provided via automatic water systems. Physical exams including body weight and complete blood counts were performed at all protocol time points. RM were sedated with ketamine HCl or Telazol for procedures, including oral and subQ vaccine administration, venipuncture, and SIV challenge.

All vaccinated RM in this study were administered a single set or 2 sets of three RhCMV/SIV vectors (68-1 backbone or 68-1 + 68-1.2 backbones), individually expressing SIV Gag, Retanef (Rev/Tat/Nef) and 5’-Pol (see below), either orally or subcutaneously at a dose of 5×10^6^ plaque-forming units per vector, with a second identical vaccination given 18 wks after primary vaccination. At the end of vaccine phase, all vaccinated and unvaccinated RM were SIV challenged by repeated (every 2-3 wks) intra-rectal administration of limiting dose (100-300 focus-forming units) SIV_mac239X_ (described below) until take of infection (onset of sustained plasma viremia and/or *de novo* development of CD4^+^ and CD8^+^ T cell responses to SIV Vif), at which time challenge was discontinued, as previously described (9, 10, 12). For *in vivo* determination of the transcriptomic response to IL-15, a cohort of five RM were treated with rRh-Het-IL-15 prepared by Drs. George Pavalakis (National Cancer Institute, USA) and Jeff Lifson (Frederick National Laboratory, USA) in a dose-escalation manner as follows: 5, 10, and 15 μg/kg at D0, D3 and D7 respectively. Whole blood was collected in PAXgene tubes at d 1, 3, 4, 7, 8, 10, 14, 17, 21, and 29 for transcriptomic analysis. RNA samples from *in vivo* rRh-Het-IL-15 treatment were processed for RNAseq transcriptomic analyses as described below.

### Ethical Statement

RM care and all experimental protocols and procedures were approved by the ONPRC Institutional Animal Care and Use Committee. The ONPRC is a Category I facility. The Laboratory Animal Care and Use Program at the ONPRC is fully accredited by the American Association for Accreditation of Laboratory Animal Care and has an approved Assurance (#A3304-01) for the care and use of animals on file with the NIH Office for Protection from Research Risks. The ONPRC adheres to national guidelines established in the Animal Welfare Act (7 U.S.C. Sections 2131–2159) and the Guide for the Care and Use of Laboratory Animals (8th Edition) as mandated by the U.S. Public Health Service Policy.

### Vectors and viruses

Construction and characterization of the 68-1 and 68-1.2 RhCMV/SIV vectors, including RhCMV/SIV_Gag_, RhCMV/SIV_Retanef(Rev/Tat/Nef)_ and RhCMV/SIV_5’-Pol (Pol-1)_ have been previously described (9, 12, 14, 15). Vector stocks were generated on telomerase-immortalized rhesus fibroblasts. SIV transgene expression was confirmed by immunoblot and all virus stocks were analyzed by next generation sequencing before *in vivo* use. Virus titers were determined by 50% tissue culture infective dose endpoint dilution assays. The pathogenic SIV challenge stocks used in these experiments were generated by expanding SIV_mac239X_ (41) in RM PBMCs and were titered using the CMMT-CD4-LTR-β-Gal sMAGI cell assay (National Institutes of Health AIDS Reagent Program).

### SIV detection assays

Plasma SIV RNA levels were determined using a gag-targeted quantitative real time/digital RT-PCR format assay, essentially as previously described, with 6 replicate reactions analyzed per extracted sample for assay thresholds of 15 SIV RNA copies/ml(6, 9, 42). Quantitative assessment of SIV DNA and RNA in cells and tissues was performed using gag targeted, nested quantitative hybrid real-time/digital RT-PCR and PCR assays, as previously described (6, 9, 42). SIV RNA or DNA copy numbers were normalized based on quantitation of a single copy rhesus genomic DNA sequence from the *CCR5* locus from the same specimen, as described, to allow normalization of SIV RNA or DNA copy numbers per 10^8^ diploid genome cell equivalents. Ten replicate reactions were performed with aliquots of extracted DNA or RNA from each sample, with two additional spiked internal control reactions performed with each sample to assess potential reaction inhibition. Samples that did not yield any positive results across the replicate reactions were reported as a value of “less than” the value that would apply for one positive reaction out of 10. Threshold sensitivities for individual specimens varied as a function of the number of cells or amount of tissue available and analyzed; for graphing consistency, values are plotted with a common nominal sensitivity threshold.

### Immunologic assays

SIV-specific CD4^+^ and CD8^+^ T cell responses were measured in peripheral blood mononuclear cells (PBMC) by flow cytometric intracellular cytokine analysis, as previously described(9, 10, 12). Briefly, individual or whole protein mixes of sequential 15-mer peptides (11 amino acid overlap) spanning the SIV_mac239_ Gag, 5’-Pol, Nef, Rev, Tat, and Vif proteins or individual SIV_mac239_ Gag supertope peptides [Gag_211-222_ (53), Gag_276-284_ (69), Gag_290-301_ (73), Gag_482-490_ (120)] were used as antigens in conjunction with anti-CD28 (CD28.2, Purified 500 ng/test: eBioscience, Custom Bulk 7014-0289-M050) and anti-CD49d stimulatory mAb (9F10, Purified 500 ng/test: eBioscience, Custom Bulk 7014-0499-M050). Mononuclear cells were incubated at 37°C with individual peptides or peptide mixes and antibodies for 1h, followed by an additional 8h incubation in the presence of Brefeldin A (5 μg ml^−1^; Sigma-Aldrich). Stimulation in the absence of peptides served as background control. After incubation, stimulated cells were stored at 4°C until staining with combinations of fluorochrome-conjugated monoclonal antibodies including: anti-CD3 (SP34-2: Alexa700; BD Biosciences, Custom Bulk 624040, PerCP-Cy5.5; BD Biosciences, Custom Bulk 624060, and Pacific Blue; BD Biosciences, Custom Bulk 624034), anti-CD4 (L200: AmCyan; BD Biosciences, Custom Bulk 658025, BV510; BD Biosciences, Custom Bulk 624340 and BUV395; BD Biosciences, Custom Bulk 624165), anti-CD8a (SK1: PerCP-eFluor710; Life Tech, Custom Bulk CUST04424), anti-TNF-α (MAB11: FITC; Life Tech, Custom Bulk CUST03355 and PE; Life Tech, Custom Bulk CUST04596), anti-IFN-γ (B27: APC; BioLegend) and anti-CD69 (FN50: PE; eBioscience, Custom Bulk CUST01282 and PE/Dazzle594; BioLegend) and for polycytokine analyses, anti-IL-2 (MQ1-17H12; PE Cy-7; Biolegend), and anti-MIP-1β (D21-1351, BV421; BD Biosciences). For analysis of memory differentiation (central- vs transitional- vs effector-memory) of SIV Gag-specific CD4^+^ and CD8^+^ T cells, PBMC were stimulated as described above, except that the CD28 co-stimulatory mAb was used as a fluorochrome conjugate to allow CD28 expression levels to be later assessed by flow cytometry, and in these experiments, cells were surface-stained after incubation for lineage markers CD3, CD4, CD8, CD95 and CCR7 (see below for mAb clones) prior to fixation/permeabilization and then intracellular staining for response markers (CD69, IFN-γ, TNF-α; note that Brefeldin A treatment preserves the pre-stimulation cell-surface expression phenotype of phenotypic markers examined in this study).

Flow cytometry analysis was preformed using an LSR-II flow cytometer (BD Biosciences). Data analysis was performed using FlowJo software (Tree Star). In all analyses, gating on the lymphocyte population was followed by the separation of the CD3^+^ T cell subset and progressive gating on CD4^+^ and CD8^+^ T cell subsets. Antigen-responding cells in both CD4^+^ and CD8^+^ T cell populations were determined by their intracellular expression of CD69 and either or both of the cytokines IFN-γ and TNF-α (or in polycytokine analyses, expression of CD69 and any combination of the cytokines: IFN-γ, TNF-α, IL-2, MIP-1β). For longitudinal immunological assessment during vaccine and challenge phases, assay limit of detection was determined, as previously described(7), with 0.05% after background subtraction being the minimum threshold used in this study. After background subtraction, the raw response frequencies above the assay limit of detection were “memory-corrected” (e.g., % responding out of the memory population), as previously described(6, 7, 9, 42), using combinations of the following fluorochrome-conjugated mAbs to define the memory vs naïve subsets: CD3 (SP34-2: Alexa700 and PerCP-Cy5.5), CD4 (L200: AmCyan and BV510), CD8a (SK-1: PerCP-eFluor710, RPA-T8: APC; BioLegend), TNF-α (MAB11; FITC), IFN-γ (B27; APC), CD69 (FN50; PE), CD28 (CD28.2; PE/Dazzle 594, BioLegend and BV510, BD Biosciences), CD95 (DX2; PE, BioLegend and PE-Cy7, BioLegend), CCR7 (15053; Biotin, R&D Systems), streptavidin (Pacific Blue, Life Tech and BV605; BD Biosciences, Custom Bulk 624342) and Ki67 (B56; FITC, BD Biosciences, Custom Bulk 624046). For memory phenotype analysis of SIV Gag-specific T cells, all CD4^+^ or CD8^+^ T cells expressing CD69 plus IFN-γ and/or TNF-α were first Boolean OR gated, and then this overall Ag-responding population was subdivided into the memory subsets of interest on the basis of surface phenotype (CCR7 vs CD28). Similarly, for polycytokine analysis of SIV Gag-specific T cells, all CD4^+^ or CD8^+^ T cells expressing CD69 plus cytokines were Boolean OR gated and polyfunctionality was delineated with any combination of the four cytokines tested (IFN-γ, TNF-α, IL-2, MIP-1β) using the Boolean AND function.

### RNA sequencing

Whole blood was collected from RM in PAXgene RNA tubes (PreAnalytiX) following the manufacturer’s procedures. (PreAnalytiX). RNA was isolated using PAXgene Blood miRNA kits (Qiagen) following the protocol provided with the kit that included an on-column DNase treatment. The quality and concentration of the recovered RNA was determined using a LabChip GXII (PerkinElmer) instrument and a ribogreen-based RNA assay, respectively. mRNA-seq libraries were constructed using Illumina TruSeq® Stranded mRNA HT kit following the manufacturer’s recommended protocol. Libraries were sequenced on an Illumina NextSeq500 sequencer using Illumina NextSeq 500/550 High Output v2 kits (150 cycles) following the manufacturer’s protocol for sample handling and loading. Sequencing run metrics were visualized for quality assurance using Illumina’s BaseSpace platform, and the quality of mRNA-seq reads were assessed using FastQC version 0.11.3 (http://www.bioinformatics.babraham.ac.uk/projects/fastqc). Both rhesus globin and ribosomal sequences were filtered via alignments with Bowtie v2.1.0 (43). Adapters were digitally removed using cutadapt, version 1.8.3: https://doi.org/10.14806/ej.17.1.200. Subsequently, a minimum of twenty million raw reads were mapped to the *Macaque mulatta* genome Mmul_1 (obtained from iGenomes: https://support.illumina.com/sequencing/sequencing_software/igenome.html) with STAR v2.4.0h1 (44) followed by HTSeq-count v0.6.1p1 (45) to generate gene counts.

### Evaluation of baseline differences

To determine whether or not a preexisting gene expression pattern was present at W0D0 that might be linked to vaccine protection, we applied hierarchical clustering using the Ward agglomerative clustering method with Euclidean distance to assess wk0, d0 time point normalized gene expression values of each animal compared to one another. Furthermore, the Pearson correlation coefficient was calculated across the wk0, d0 data set, showing that all animals of the training set exhibited wk0, d0 baseline gene expression signatures similar to one another independently of vaccine protection outcome.

### Preparation for analysis of differential expression

Based on the raw read counts, outlier samples and genes with a maximum expression across all samples below 100 counts were removed. Using R (v 3.6.0)/Bioconductor(v 3.9), counts were then transformed into counts per million using the voom function(46) in the R library *limma* (47) with a smoothing window of 0.1. CPMs were normalized using the quantile method. Differential expression was performed using the *limma* package in R/Bioconductor (48). Additional graphics packages were utilized for the visualization of numbers of DE genes (*ggplot2;* https://ggplot2.tidyverse.org) and heat maps (*gplots;* https://www.rdocumentation.org/packages/gplots/versions/3.1.0) using wk0, d0 as the common baseline comparator for each animal in the training set and the validation set cohorts.

### Differential expression analysis

To determine the list of significantly differentially expressed genes in the RhCMV/SIV vector-vaccinated cohort as well as in the rRh-Het-IL-15 experiment, we used the *lmFit* function in the R library *limma* (47). Genes with a false discovery rate (FDR)-adjusted p-value ≤ 0.05 and absolute log_2_ fold change (compared to baseline) above 1.5 were defined as significantly differentially expressed (DE). To define gene correlates of vaccine protection we determined the set of genes for which the baseline-subtracted expression values significantly differed between protected and non-protected outcome groups of the training set cohort (DDE). Genes with FDR-adjusted p ≤ 0.05 and absolute log2(FC) (across protection groups) above 1.5 were defined as significantly differentially DE (DDE). These genes were identified using the interaction effect between time point (compared to baseline) and vaccine protection.

### Principle component analysis (PCA)

PCA was performed on the per-timepoint mean log2(FC) values, using all expressed genes and averaged over the RMs within each treatment and outcome group. We used the R function PCA in the *FactoMineR* library, with variance scaling enabled and keeping 7 dimensions in the output. For each of the top 7 dimensions, we included the genes most highly correlated with the PC. The number of genes selected from each dimension was the dimension’s percentage explained variance times 2.5 (an arbitrary value selected to balance figure size with information content).

### Exploratory pathway analyses

Enrichment tests and network analyses for pathways and upstream regulators were performed using Ingenuity Pathway Analysis (49) (Sept 2019 version for Fig. 2; March 2020 version for Fig3). IL-15-regulated networks were identified using GeneMANIA(50). Co-expression analyses were performed only on DE genes. We conducted clustering analysis using Ward clustering and Euclidean distance on the union of log2(FC) values using the *WGCNA*, *heatmap.2*, and *EdgeR* Bioconductor packages in R(51–53).

### Hierarchical cluster analysis and heatmap generation

For correlation analysis of the fold-change values displayed in heatmaps, we used midweight bicorrelation, a modified version of Pearson correlation (bicor function in the R library *WGCNA*), and complete linkage clustering. The clusters were defined using the cutree function (R library *stats*) (54). All trees were cut with height 1.4, which corresponds to a midweight bicorrelation coefficent of −0.4 because the analysis is constructed by shifting the correlations to the range (0,2), resulting in four clusters in both instances. The heatmaps were drawn using the heatmap.2 function (R library *gplots*).

### Permutation testing

To formally test whether the DDE and pathway-linked gene signatures significantly differentiate protected vs. unprotected RMs over the course of study, we devised and followed a formal statistical analysis plan. Briefly, we defined a test statistic aggregating over genes and time and compared this to a null distribution that controls for the observed data (including all correlations across genes, which is ignored in the primary linear modeling analysis described above). In this procedure we first calculate, for each gene, the absolute value of the mean over time of the difference in log2(FC) across protection groups. For the DDE analysis, the primary test statistic is the sum of the above value across all significantly DDE genes. We then compare this value to its empirical null distribution, approximated by sampling protection outcomes within each treatment group over 5000 permutations sampled with replacement. Note that for each permutation, the list of significantly DDE genes is allowed to change (the value of the test statistic was set to zero for permutations in which no genes were significantly DDE). We then repeated this analysis, where instead of the DDE gene list, we used the gene lists related to the IL-15 response, TCR signaling pathway, TLR signaling pathway, and Inflammasome pathway. These gene lists are fixed and do not vary across permutations; otherwise, all other aspects of the fixed-list analyses were identical to the DDE permutation analysis. Unadjusted p-values are the proportion of the 5000 permutations ≥ the observed test statistic (a one-sided test; the statistic is an absolute value, so it is always positive). To evaluate the hypothesis that the IL-15 response signature component of DDE genes is expressed among protected RM more than non-protected RM, we devised a consistency test statistic that directly addresses the hypothesis that protected RM have a response to vaccination consistent with a response to IL-15 administration. This test statistic compares across protection categories a summary measure indicating the extent to which the gene response to RhCMV/SIV vaccination is consistent with the response to IL-15 administration (up-regulated genes going up, down-regulated genes going down), by measuring the difference across two gene lists of the sum of the average log2(FC) over time: the total among those genes up-regulated by IL-15 administration minus the total among those down-regulated by it. For these analyses, instead of sampling from all possible permutations with replacement, we evaluated the support of the null hypothesis exhaustively resulting in an exact test (with 5005 possible configurations in evaluating the validation cohort, and 6435 when evaluating the subQ cohort). Unadjusted p-values for these tests are the proportion of the null distribution ≥ the observed test statistic (testing a one-sided hypothesis).

### Additional statistical analyses

Pre-vaccination transcriptome correlation analysis was conducted using Pearson correlation. Viral load and immunologic data are presented as boxplots with jittered points and a box from 1st to 3rd quartiles (IQR) and a line at the median, with whiskers extending to the farthest data point within 1.5×IQR above and below the box. Analyses of longitudinal ICS data were performed by calculating the per-RM average T-cell response over three periods: post-prime peak (2-6 weeks), post-boost peak (20-24 weeks), and plateau (61-90 weeks), and comparing these values between RMs receiving subQ vs. oral vaccine using the nonparametric Wilcoxon rank-sum test. All P*-*values are based on two-sided tests and unadjusted except where noted. Adjusted P*-*values were computed using the Holm procedure for family-wise error rate control.

## Data and code availability

Transcriptomics data sets are deposited at the Gene Expression Omnibus https://www.ncbi.nlm.nih.gov/geo/ under accession number GSE160562. The R markdown code applied to these analyses can be accessed at https://github.com/galelab/GaleGEAnalysis and at https://github.com/komorowskilab/R.ROSETTA.

### Links to software

STAR aligner: https://github.com/alexdobin/STAR

Bowtie2: http://bowtie-bio.sourceforge.net/bowtie2/index.shtml

HTseq: https://github.com/simon-anders/htseq

FASTQC: https://www.bioinformatics.babraham.ac.uk/projects/fastqc/Cutadapt: https://github.com/marcelm/cutadapt

Venny: http://bioinfogp.cnb.csic.es/tools/venny/index.html

Ingenuity Pathway Analysis: https://digitalinsights.qiagen.com/products-overview/discovery-insights-portfolio/analysis-and-visualization/qiagen-ipa/Genemania: https://genemania.org/

## Author contributions

SGH planned and performed animal experiments and immunologic assays, assisted by CMH, DM, KR, ANS, JCF, EA, and RMG. MKA supervised animal procedures and care. JDL planned and supervised SIV quantification by PCR/RT-PCR assisted by KO, RS, RF, and WJB. GNP, BKF, and JDL prepared and provided rRh-Het-IL-15 and developed methodology for use. YF and BER performed the *in vivo* analysis of rRh-Het-IL-15 administration to RM. LL supervised sample intake, processing, and assignment for RNAseq. ES, JC, and IG conducted RNA processing, library construction, sequencing, and computational quality control of sequence reads. FB conducted bioinformatics analyses, including linear modeling. JK developed linear modeling applications. CD, RRG, XP, LW, DN, SF, and RPS conducted specific bioinformatics analyses. PTE planned, conducted and supervised statistical analyses, assisted by EW, JS, and WS. LJP conceived the RhCMV vector strategy, supervised all RM experiments and immunologic analyses, analyzed and interpreted data. MG led the RNA sequencing and bioinformatics analyses. LJP and MG cowrote the manuscript. Correspondence and request for materials should be addressed to either MG or LJP.

## Acknowledgments

This work was supported by the National Institute of Allergy and Infectious Diseases (NIAID) contract HHSN272201800008C, and grants P30 AI027757-31 and P51 OD010425 (MG); grants P01 AI094417, U19 AI128741, UM1 AI124377, and R37 AI054292 (LJP), the Oregon National Primate Research Center Core grant from the National Institutes of Health, Office of the Director (P51 OD011092); by the Intramural Research Program of the National Cancer Institute (GNP and BKF), and by contracts from the National Cancer Institute (# HHSN261200800001E; JDL). The authors thank C. Bergamaschi, J. Bear, E. Chertova, R. Sowder, D. Roser, and J. Bess, Jr., for purification and biochemical characterization of the rRh-Het-IL-15 and information on its use. OHSU, LJP and SGH have a substantial financial interest in Vir Biotechnology, Inc., a company that may have a commercial interest in the results of this research and technology. LJP and SGH are also consultants to Vir Biotechnology, Inc. Other authors have no potential conflicts of interest.

## Abbreviations

Bo: boost
DE: differentially expressed
DDE: differential expression of DE genes (protected vs. non-protected)
EM: effector-memory
FC: fold-change
FDR: false discovery rate
ICS: intracellular cytokine staining
IRF: interferon regulatory factor
ISGs: interferon stimulated genes
ONPRC: Oregon National Primate Research Center
PI3K: phosphatidylinositol responsive kinase
Pr: prime
preCh: pre-challenge
PC: principal component
PCA: principal component analysis
RhCMV: Rhesus Cytomegalovirus
RMs: rhesus macaques
SIV: Simian Immunodeficiency Virus
subQ: subcutaneous

## Supporting Information

### Supplemental Tables

Table S1: SubQ and oral cohort fold-changes (from baseline; FC), unadjusted P-values, and adjusted P-values of DE genes of 68-1 oral and subQ cohort RMs vaccinated with RhCMV/SIV.

Table S2: Estimates of differences in DE across protection groups for DDE genes of 68-1 oral and subQ cohort RMs vaccinated with RhCMV/SIV, with unadjusted and adjusted P-values.

Table S3: SubQ and oral cohort fold-changes (from baseline), unadjusted P-values, and adjusted P-values of IL-15 response genes for all ds (Sheet 1) and 1d after rRh-Het-IL-15 administration (Sheet 2).

Table S4: SubQ and oral cohort fold-changes (from baseline), unadjusted P-values, and adjusted P-values of DDE genes in 68-1 oral and subQ cohort RMs that are also IL-15 response genes.

Table S5: Validation cohort fold-changes (from baseline), unadjusted p-values, and adjusted P-values of validation cohort DE genes.

Table S6: Validation cohort fold-changes (from baseline), unadjusted p-values, and adjusted P-values of DDE genes in 68-1 oral and subQ cohort RMs that are also IL-15 response genes.

### Supplemental Figures

Figure S1. Immunogenicity of 68-1 RhCMV/SIV vectors in subcutaneously vs. orally vaccinated RM.

Figure S2. The magnitude and phenotype of SIV-specific CD4^+^ and CD8^+^ T cell responses in blood do not predict 68-1 RhCMV/SIV vector efficacy.

